# Mutations in CmVPS41 controlling resistance to *Cucumber Mosaic Virus* display specific subcellular localizations

**DOI:** 10.1101/2022.08.30.505877

**Authors:** Núria Real, Irene Villar, Irene Serrano, Cèlia Guiu-Aragonés, Ana Montserrat Martín-Hernández

## Abstract

Resistance to *Cucumber mosaic virus* (CMV) in melon has been described in several exotic accessions. It is controlled by a recessive resistance gene, *cmv1*, which encodes a Vacuolar Protein Sorting 41 (CmVPS41). *Cmv1* prevents systemic infection by restricting the virus to the bundle sheath cells, preventing viral phloem entry. CmVPS41 from different resistant accessions carried two causal mutations, either a G85E change, found in Pat-81 and Freeman’s Cucumber, or L348R found in PI161375, cultivar Songwhan Charmi (SC). The analysis of the subcellular localization of CmVPS41 in *N. benthamiana* has revealed differential structures in resistant and susceptible accessions. Susceptible accessions showed nuclear and membrane spots and many transvacuolar strands, whereas the resistant accessions showed many intravacuolar invaginations. These specific structures colocalize with late endosomes. Artificial CmVPS41 carrying individual mutations causing resistance in the genetic background of CmVPS41 from the susceptible variety Piel de Sapo (PS), revealed that the structure most correlated with resistance was the absence of transvacuolar strands. Co-expression of CmVPS41 with the viral MPs, the determinant of virulence, did not change these localizations; however, infiltration of CmVPS41 from either SC or PS accessions in CMV-infected *N. benthamiana* leaves showed a localization pattern closer to each other, with up to 30% cells showing some membrane spots in the CmVPS41SC and fewer transvacuolar strands (from a mean of 4 to 1-2) with CmVPS41PS. Our results suggest that the distribution of CmVPS41PS in late endosomes includes transvacuolar strands that facilitate CMV infection and that CmVPS41 is re-localized during viral infection.

## Introduction

When viruses enter the plant, they must replicate, move cell-to-cell generally through plasmodesmata, up to the veins, surrounded by the bundle sheath cells (BS), and invade the phloem cells, namely vascular parenchyma cells (VP) and companion cells (CC) to finally invade the whole plant, producing a systemic infection (Hipper et al., 2013). Since viruses have a small and compact genome, encoding only a few genes, they must request the participation of many host factors to complete their cycle. Mutations in those host factors develop a loss of susceptibility that can either limit or prevent viral infection, thus, becoming resistance genes that are recessively inherited (Hashimoto et al., 2016). Therefore, understanding how viral proteins and virions interact with host factors is key to prevent viral diseases.

*Cucumber Mosaic Virus* (CMV) is a positive-stranded RNA virus able to infect over 1200 plant species, including members of three important crop families, Solanaceae, Cruciferae and Cucurbitaceae (Edwardson and Christie, 1991). CMV genome has three genomic and two subgenomic RNAs that encode five viral proteins. CMV strains are divided in two subgroups, I (SG I) and II (SG II) which share 70% of their sequence and present differences in their serological and chemical properties (Roossinck, 2001). In melon few sources of resistance to CMV have been identified (Karchi et al., 1975; Pascual et al., 2019; Martín-Hernández and Picó, 2021), among them the Korean cultivar ‘Songwhan Charmi’, PI 161375 (from now on SC). SC shows an oligogenic and recessive resistance (Karchi et al., 1975), with a major gene, *cmv1*, conferring resistance to strains of SG II (Essafi et al., 2009; Guiu-Aragonés et al., 2015), and at least two other Quantitative Trait Loci (QTLs) which, together with *cmv1* confer resistance to SG I strains (Guiu-Aragonés et al., 2014). The resistance conferred by *cmv1* is manifested as a restriction to phloem entry, since in the plants carrying this gene, strains of SG II (such as CMV-LS) can replicate and move cell-to-cell in the mesophyll up to the BS cells but are restricted in these cells and cannot move to the phloem cells. However, strains of SG I (such as CMV-FNY) can overcome this restriction and invade the phloem (Guiu-Aragonés et al., 2016). The viral factor determining this ability is the movement protein (MP), since a viral clone from CMV-LS carrying the MP from FNY can invade the phloem of the plant carrying the gene *cmv1* (Guiu-Aragonés et al., 2015).

Map-based cloning in melon demonstrated that *cmv1* encodes a Vacuolar Protein Sorting 41 (CmVPS41) (Giner et al., 2017), a protein involved in the intracellular vesicle transport of cargo proteins from the late Golgi to the vacuole as part of the “homotypic fusion and vacuole protein sorting” (HOPS) complex, both via endosomes and also through vesicles, via the AP-3 pathway (Balderhaar and Ungermann, 2013; Schoppe et al., 2020). HOPS is a complex of six subunits, four of them shared (VPS11, VPS33, VPS16 and VPS18) with CORVET, another complex involved in endosome life cycle. The remaining two, VPS39 and VPS41 are required for the tethering function, being VPS41 the effector subunit of HOPS to promote vacuole fusion (Price et al., 2000). VPS41 is localized in late endosomes in Arabidopsis (Brillada et al., 2018). In yeast, VPS41p participates in the membrane fusion of cargo proteins between late endosomes and lysosomes (Rehling et al., 1999). In mammals self-assembly of VPS41 is required for the biogenesis of the secretory pathway (Asensio et al., 2013) and mutations in this protein are related to neurological disorders and abnormal membrane trafficking (Sanderson et al., 2021) associated with lysosomal abnormalities (Steel et al, 2020). Likewise, deletions in VPS41 in pancreatic B cells cause defects in insulin secretion (Burns et al, 2021). In fungi, VPS41 also localizes to endosomes and vacuole membrane, and it is essential for plant infection and fungal development (Li et al., 2018). In *Arabidopsis thaliana*, AtVPS41 controls pollen tube-stigma interaction and is found in prevacuolar compartments and in the tonoplast, where it is required for the late stage of the endocytic pathway (Hao et al., 2016). It is highly expressed in anthers and petals, followed by roots and cotyledons (Klepikova et al., 2016). This high expression in anthers fits with the inability to develop pollen tubes in some mutants, leading to male sterility (Hao et al., 2016). Additionally to its localization to the tonoplast, in root cells AtVPS41 is also located to condensates, that are essential for developmental regulation (Jiang et al., 2022). T-DNA insertion mutants in Arabidopsis seem to be lethal, revealing the fundamental role of VPS41, whereas *zip-2*, a single mutant in AtVPS41, shows no obvious phenotype (Niihama et al., 2009). Likewise, point mutations observed in melon CmVPS41 show no phenotype except impairing CMV phloem entry, suggesting that the main function of VPS41 has not been altered with these mutations (Giner et al., 2017; Pascual et al., 2019). VPS41 has also been related to Ebola and Marburg flaviviruses infection in humans (Carette et al., 2011).

Expression of CmVPS41PS transgene in the melon accession SC can complement CMV phloem entry, allowing a systemic infection (Giner et al., 2017). Although CmVPS41 is a general gatekeeper in many melon genotypes for viral phloem entry and determines the resistance against CMV-LS, its mechanism of action is yet to be characterized. Here we have investigated the localization pattern of CmVPS41 from both, susceptible and resistant melon genotypes, and the localization of CmVPS41 carrying only the mutations causing the resistance from some resistant accessions. We have also investigated their response to the presence of the viral MP and to the whole virus during the infection.

## Material and Methods

### Plant material, viral strains, and yeast strains

*Cucumis melo L*. genotypes, Piel de Sapo (PS), Songwhan Charmi (SC), Freeman’s Cucumber (FC), Pat-81 and Cabo Verde (CV), with different susceptibilities to CMV were used to clone their corresponding CmVPS41 genes (Supplemental table 1). The list of CmVPS41 included two chimerical constructs with CmVPS41 genotype from PS carrying the identified causal mutations: (i) causal mutation present in cultivar SC (L348R) (Giner et al., 2017) and (ii) causal mutation found in cultivars FC and Pat 81 (G85E) (Pascual et al., 2019). *Cucumis melo L*. seeds were pre-germinated by soaking them in water overnight, and then maintained for 2-6 days in neutral day at 28 °C. Seedlings were grown in growth chambers SANYO MLR-350H in long-day conditions consisting of 22°C for 16 h with 5000 lux of light and 18 °C for 8 h in the dark. For agroinfiltration, *N. benthamiana* plants were grown in the greenhouse in long-day conditions consisting of 24-28 °C with 5000 lux of light for 16 h and 22-24 °C 8 h in the dark.

CMV strains CMV-LS and CMV-FNY were used for CMV infection assay. *S. cereviseae* strains Y2HGold and Y187 were used for Yeast Two Hybrid assay (Takara Bio, Mountain View, USA).

### Plasmid construction

For co-localization experiments with CmVPS41s, total RNA extractions from the different melon accessions were performed using TriReagent (SIGMA-ALDRICH, St Louis, MO, USA) following the manufacturer’s protocol. 200 ng of total RNA were used to synthesize cDNA using oligo (dT)_12-18_ primer (Invitrogen by Thermo Fisher Scientific, Vilnius, Lithuania) and PrimeScript (Takara Bio, Dalian, China), according to manufacturer’s instructions. For cloning the complete CmVPS41 genes, cDNA was PCR-amplified using the PrimeSTAR® GXL DNA Polymerase (Takara Bio, Dalian, China) and the primers annealing at the ends of the gene and carrying the attB sequences (Supplemental table 1) and cloned into the pBSDONR P1-P4 (Gu and Innes, 2011) by Gateway BP reaction (MultiSite Gateway® Pro from Invitrogen by Thermo Fisher Scientific, Vilnius, Lithuania). For CmVPS41 clones carrying the causal mutations, CmVPS41 G85E and L348R constructs were generated from CmVPS41 pBSDONR P1-P4 VPS41PS and VPS41SC (for L348R) or VPS41PS and VPS41FC (for G85E) constructs using a combination of specific primers (Supplemental table 1) to transfer the fragments containing the causal mutations to the VPS41PS background using Gibson Assembly technology (GeneArt® Seamless PLUS Cloning and Assembly Kit, Invitrogen Corporations, Carlsbad, CA, USA). eGFP (Cormack et al., 1996) and RFP (Campbell et al., 2002) were cloned into pBSDONR P4r-P2 for C-terminal fusion using specific primers (***Error! Reference source not found***.). The P1-P4 clones were mixed with the corresponding P4r-P2 and the dexamethasone-inducible destination vector pBAV154 (Vinatzer et al., 2006) in a three-way Gateway LR reaction (MultiSite Gateway® Pro from Invitrogen by Thermo Fisher Scientific, Vilnius, Lithuania).

For MP localization experiments, the whole MP gene of both strains was PCR amplified using primers that generate *BamHI* and *XhoI* at 5’ and 3’ end of gene respectively. PCR products were cloned into GATEWAY® pENTR− 3C (InVitrogen Corporations, Carlsbad, CA, USA) at *BamHI-XhoI* sites. To express C-terminally tagged fluorescent protein fusion of MP:GFP, both pENTR-MP-LS and pENTR-MP-FNY were recombined with destination vector pH7WGF2 (Karimi et al., 2002), using LR clonase mix (Invitrogen Corporations, Carlsbad, CA, USA) according to the manufacturer’s instructions. pDLP1:GFP clone was kindly provided by Prof. Andy Maule (John Innes Centre) For Bimolecular Fluorescence Complementation assay (BIFC), MPs and CmVPS41s coding sequences were PCR-amplified using specific primers (Supplemental table 1) and cloned into pDONR P1-P4 as described above, and introduced into the expression vector pBAV154 as a C-terminal fusion with either partial N-terminal YFP (YN) (for CmVPS41s) or partial C-terminal YFP (YC) (for CMV MPs) (Supplemental table 2) using Gateway technology, as previously described.

For Yeast Two Hybrid experiments, coding sequences of CMV MPs and CmVPS41s were PCR amplified using specific primers (Supplemental table 1) and cloned into plasmid pENTR/D-TOPO (Thermo Scientific by Thermo Fisher Scientific, Vilnius, Lithuania). Both MP genes were cloned into the pGBKT7 vector at the *BamHI* and *EcoRI* sites using T4 DNA ligase (Thermo Scientific by Thermo Fisher Scientific, Vilnius, Lithuania) following manufacturer’s instructions. CmVPS41s were cloned into pGADT7 vector at the *SalI* and *EcoRI* sites using T4 DNA ligase (Thermo Scientific by Thermo Fisher Scientific, Vilnius, Lithuania).

For co-localization of CmVPS41 with organelle markers, *Arabidopsis thaliana* genes were used (Supplemental table 2). Markers for Endoplasmic reticulum (35S:HDEL-mCherry), Golgi Apparatus (35S:MAN49-mCherry), Tonoplast (35S:γ-TIP-mCherry) and late endosome (ARA6-mCherry) were from (Serrano et al., 2016). Plasma membrane marker (35S:Remorin-mCherry) was nicely provided by Dr. Núria Sanchez Coll (CSIC, CRAG, Barcelona, Spain) (Marín et al., 2012).

All constructs were verified by sequencing with Sanger method using an ABI 3730 DNA Analyzer (Applied Biosystems) for capillary electrophoresis and fluorescent dye terminator detection. Correct insertion and orientation of all constructs were verified with Sequencher® version 5.0 sequence analysis software (Gene Codes Corporation, Ann Arbor, MI, USA (http://www.genecodes.com). Correct plasmids were transformed into *A. tumefaciens* GV3101.

### Transient expression in *Nicotiana benthamiana*

*A. tumefaciens* cultures carrying the corresponding plasmid were incubated at 28ºC with their corresponding antibiotics for 24-48 h and bacterial pellet was collected by centrifugation and resuspended in Induction Buffer (1M MgCl_2_ and 0.15 M acetosyringone) to 0.4 final OD_600_, except for the 35S:Remorin 1.3-mCherry experiments, where 0.2 final OD_600_ was used. Bacterial culture was induced for 2 h in the dark. For agroinfiltration of more than one plasmid, suspensions were mixed in equal ratio and were infiltrated using a needleless syringe into the abaxial side of expanding leaves of 2–3-week-old *N. benthamiana* plants. For dexamethasone-inducible pBAV154-derived constructs, 24 h post infiltration, expression was induced applying 50 μM dexamethasone solution with a brush in the adaxial part of the infiltrated leaf (Sigma, St. Louis, USA). Expression of fluorescence was observed at 20 h after dexamethasone induction in pBAV154-derived vectors and 48 h after agroinfiltration in vectors with 35S promoter.

### Yeast Two Hybrid assays

Constructs for Yeast Two Hybrid (Y2H) were freshly transformed into *S. cereviseae* strains before every one-to-one Y2H assay. pGBKT7-MPs, positive control (pGBKT7-53) and negative control bait plasmids (pGBKT7-Lam) were transformed into yeast strain Y2HGold (Takara Bio, Mountain View, USA) while pGADT7-CmVPS41s and pGADT7-T control prey plasmids were transformed into yeast strain Y187 following the Yeastmaker Yeast Transformation System 2 instructions (Clontech by Takara Bio, Mountain View, USA). The transformed cells were grown either in SD-Trp agar plates for Y2HGold or in SD-Leu agar plates for Y187 at 30 ºC for 3-5 days. Matings between the Y2HGold and Y187 strains carrying the appropriate constructs were performed in 0.5 mL of 2X YPDA in the presence of the corresponding antibiotics following manufacturer’s instructions (Clontech’s Matchmaker Gold Yeast Two-Hybrid System, Clontech by Takara Bio, Mountain View, USA). Yeast cells were cultured at 30 ºC for 24 h and plated in SD-Trp/-Leu/X-alpha-Gal agar plates. After 3-5 days blue or white colonies appeared and were transferred in a more restrictive media (SD-Trp/-Leu/X-alpha-Gal/Aureobasidin A agar plates).

After 5-7 days blue colonies (or in its defect white colonies) were selected and plated in the most restrictive media (SD-Ade/-His/-Trp/-Leu/X-α-Gal/AbA). True interaction was assumed when strong blue colonies were able to grow in the more restrictive media.

### CMV inoculations

Viral inocula were freshly prepared from infected zucchini squash Chapin F1 (*Cucurbita pepo L*.) (Semillas Fitó SA, Barcelona, Spain). Sap was rub-inoculated in first and second true leaves of 2-week-old *N. benthamiana* plants.

### Virus detection

First true leaf of two *N. benthamiana* plants was inoculated with either CMV-LS or CMV-FNY. After detection of the symptoms, one leaf of each plant was collected at two- and three-weeks post-inoculation. RNA was isolated, and the presence of the virus in the leaves was tested by reverse transcription PCR as described (Guiu-Aragonés et al., 2015). CMV-LS, specific primer LS1-1400R, LS2-1400R, LS3-1400R, PCR1-1400R, PCR2-1400R and PCR2-1400R were used for Reverse transcription reaction. The same primers, together with LS1-900F, LS2-900F, LS3-900F, FNY1-900F, FNY2-900F, FNY3-900F, were used to amplify a 500-bp fragment of each RNA from the viral genome (Supplemental table S1).

### Confocal Laser Scanning Microscopy

For Bimolecular fluorescence complementation (BiFC) and co-localization experiments of plasma membrane or endoplasmic reticulum with CmVPS41 proteins, images were collected on a Leica TCS-SP5 II confocal microscope (Leica Microsystems, Exton, PA USA) using a 63x water immersion objective NA 1.2, zoom 1.6. In BiFC images YFP was excited with the blue argon ion laser (514 nm), and emitted light was collected between 530 nm and 630 nm using a HyD detector. For co-localization, eGFP was excited with the blue argon ion laser (488 nm), and emitted light was collected between 495–535 nm. mCherry and RFP were excited with an orange HeNe laser (594 nm), and emitted light was collected between 570–660 nm. Chloroplasts were excited with the blue argon laser (488 nm), and emitted light was collected at 650-750 nm. eGFP and chloroplasts signals were collected separately from the mCherry or RFP signals and later superimposed. All images were processed using Fiji imaging software (version 1.52i). Fiji co-localization tool “Co-localization Threshold” was used to calculate the Pearson coefficient of co-localization and to create a colocalized Pixel Map for each combination of CmVPS41 plus organelle marker. To observe CmVPS41 transvacuolar strands IMARIS software (Bitplane AG, Zurich, Switzerland) was used to perform 3D reconstructions of the Z-stacks of confocal images and capture snapshots.

In all other agroinfiltration experiments, images were collected on Olympus FV1000 confocal laser scanning microscope (Olympus, Tokyo, Japan). During scanning, we used a quadruple-dichroic mirror (DM 488/559). For visualization of eGFP, RFP, mCherry and chloroplasts the same emission and collection windows than for the Leica Microscope were used. The images with co-localization were studied by sequential excitation with each laser separately to avoid crossed fluorescence in the red channel. Images were processed using Olympus FV1000 software (version 04.02). Fiji (1.52i) was used to set the scale bar and calculate the Pearson coefficient for co-localization experiments with CmVPS41 and organelle markers.

### Western blot

For determining the right expression of CmVPS41s proteins, 30 mg of frozen leaf powder per agroinfiltration were used. Protein extraction was carried out directly in 100 μl SDS sample buffer [2% (w/v) SDS, 62.5 mM Tris–HCl (pH 6.8), 10% (v/v) glycerol and 0.007% (w/v) bromophenol blue] including 50 mM DTT and protease inhibitor (Roche) and then boiled (95ºC) for 5 min. 30 μl were loaded and resolved on 6% sodium dodecyl sulfate (SDS) gels and transferred onto nitrocellulose membranes with Mini-PROTEAN III and Mini Trans-Blot cells (Bio-Rad, Hercules, CA, USA), respectively, following conventional protocols. Uniform protein loading and transfer efficiency were verified by Ponceau staining of membranes. Primary antibody was anti-green fluorescent protein (mouse anti-GFP, Abcam, Cambridge, UK, Ab291-50, 1:1000). The secondary antibody was anti-mouse IgG horseradish peroxidase conjugate (Sigma, St. Louis, MO, USA). Immunoreactive proteins were detected by chemiluminescence using the SuperSignal West Femto kit (Pierce, Thermo Fisher Scientific, Waltham, MA, USA).

## Results

### CmVPS41 from both PS and SC associate in vivo with CMV-FNY Movement Protein

Given that CMV MP is the determinant of virulence that communicates with CmVPS41, we investigated if there was interaction between these two proteins. For Y2H experiments, yeast clones carrying either MP-FNY or MP-LS were tested against those carrying either CmVPS41PS or CmVPS41SC. After mating, the resulting colonies were as white as the negative control, while the positive control turned blue. This indicated that none of the MPs interacted with none of the CmVPS41s (Figure 1A). This experiment was repeated twice, getting the same result. To confirm this result *in vivo*, we performed BiFC experiments using the constructs VPS41s-YN and MPs-YC, where the CmVPS41 from either PS or SC carried the N-terminal part of YFP, and the MP from either CMV-FNY or CMV-LS carried the C-terminal part of YFP. Then, co-agroinfiltration of the VPS41s-YN and MPs-YC BiFC constructs showed that MP-FNY was able to interact both with CmVPS41PS and CmVPS41SC in a pattern compatible with their localization at the plasmodesmata, whereas MP-LS was unable to interact with any CmVPS41. CmL-ascorbate-oxidase was used as positive control for interaction with CMV MP (Figure 1C). Negative controls were the same proteins alone (Supplemental Figure S1). To confirm that CMV MPs localize at the PDs, we investigated the cellular localization of the CMV MPs alone. Co-agroinfiltration of a 35S:MP-FNY-GFP or 35S:MP-LS-GFP constructs with the PD marker PDLP1 (Plasmodesmata-located protein 1) (Amari et al., 2010) tagged with RFP, showed that, as expected, both MPs co-localize with PDLP1 at the PDs (Figure 1B). Therefore, the interaction between VPS41s and MP-FNY was taking place at or near the PDs. VPS41s-YN would also localize at the same localizations shown in figure 3 (tonoplast, late endosome, plasma membrane) but it would only be visualized where CMV MP was present. This experiment does not preclude the possibility that MP FNY re-localizes at least part of CmVPS41 to PDs Altogether, these experiments indicated that, the only possible interaction was between the CmVPS41PS or CmVPS41SC with MP-FNY.

**Figure 1.**
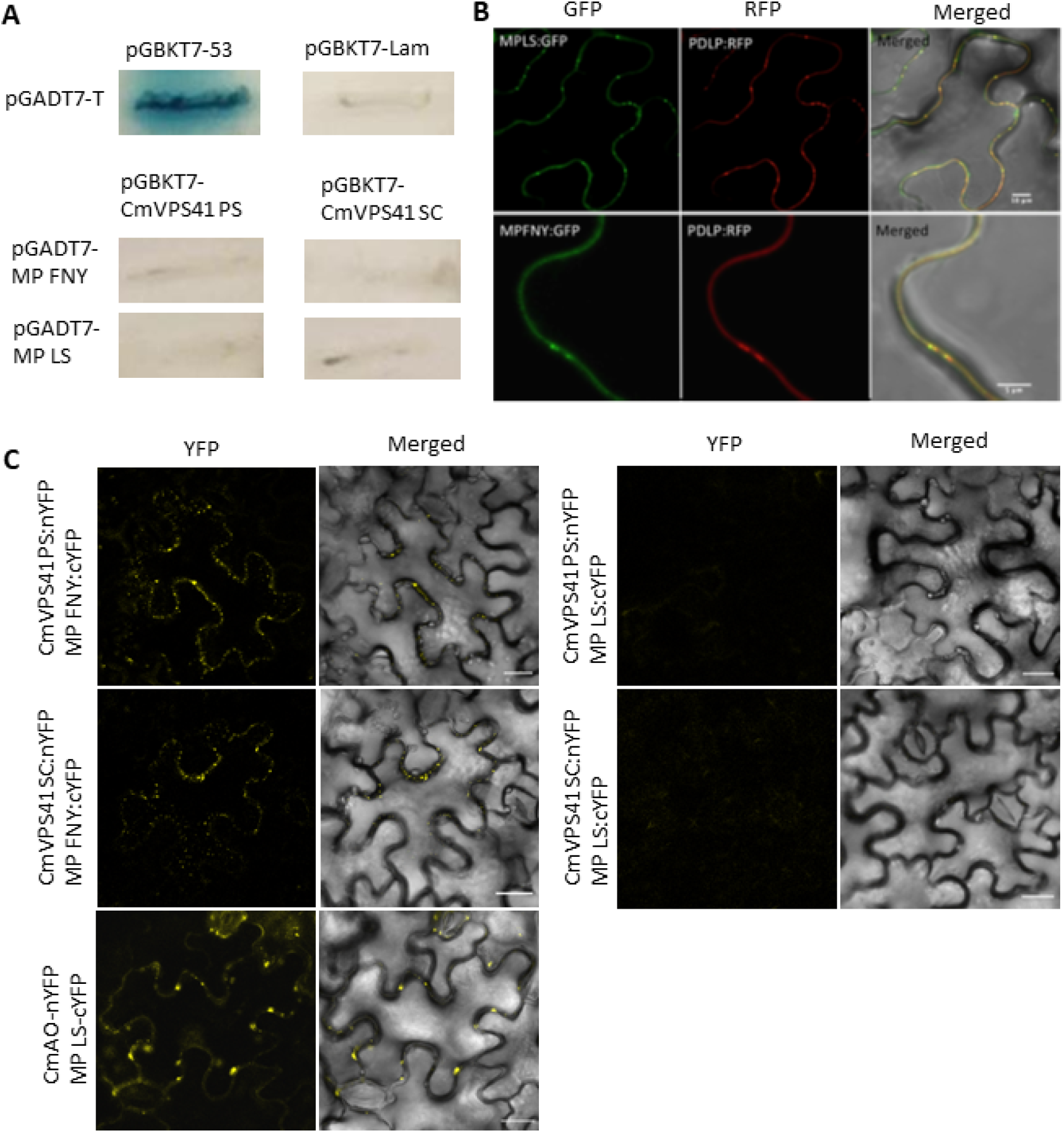
Interaction between CmVPS41s and CMV MPs. **A**. Yeast-Two-Hybrid (Y2H) between CmVPS41 and CMV MPs in SD-Trp/-Leu/X-alpha-Gal/AbA/-His/-Ade agar plates. Each cell shows the results ofY2H interaction combination of prey in vector pGADT7 per bait in vector pGBKT7. Growth of a strong or light blue colony indicates interaction between prey and bait, whilst absence of growth or white colonies indicates no interaction. Controls are pGADT7-T (prey) in combination with either bait pGBKT7-53 (positive interaction) or bait pGBKT7-Lam (no interaction). **B**. Co-localization of CMV MPs (MP-FNY-GFP or MP-LS-GFP) with plasmodesmata marker (PDLP1-RFP). Co-localization of MPs and PDLP can be observed as yellow colour. C. In planta BIFC assay between CmVPS41s and CMV MPs. CmAO-nYFP, L-ascorbate oxidase used as positive MP interacting control. ‘Merged’: YFP and bright field channel together. BIFC scale bars correspond to 20 pm length.

### Localization pattern of CmVPS41 from resistant and susceptible melon genotypes is different

To analyze the subcellular localization of CmVPS41s, the genes from both, the susceptible melon genotype PS and the resistant SC, tagged with eGFP, were expressed under a Dexamethasone-inducible promoter in *N. benthamiana* epidermal cells. We observed that 20 hours after Dexamethasone application, both variants seem to localize to the cytoplasm and the nucleus. However, there were some differences in the expression pattern of both CmVPS41 variants. CmVPS41PS expression was very strong as spots in the plasma membrane or in the tonoplast and presented speckles in both, the nuclear membrane, and the nucleus. Also, there were several transvacuolar strands in most cells (Figure 2A, 1 and 2). However, in the cells expressing the resistant allele, CmVPS41SC, there were no membrane spots, with few or no speckles, the expression inside the nucleus and cytoplasm was smooth, there were no transvacuolar strands and there were many tonoplast invaginations towards the vacuole (Figure 2A, 3 and 4). These localization patterns were clearly different from those shown when expressing eGFP alone (Supplemental Figure S2). Western blot analyses showed that both CmVPS41 proteins had been correctly expressed (Supplemental Figure S3 and S4, lanes 1 and 2). A quantification of the cells carrying these distinctive structures showed significant differences among the susceptible PS and the resistant SC variants (Figure 2B). More than 90% of cells expressing CmVPS41PS showed nuclear speckles, whereas in only 30% of cells expressing CmVPS41SC some speckles could be found. The difference in membrane or tonoplast spots was even more evident, since almost none of the CmVPS41SC-expressing cells showed them, whereas nearly all cells expressing CmVPS41PS presented them. The differences in transvacuolar strands were also significant, with most CmVPS41SC cells presenting one or none and most CmVPS41PS presenting a mean of four strands. Last, a mean of 75% of VPS41SC expressing cells showed tonoplast invaginations, whereas those structures were almost absent from CmVPS41PS-expressing cells. Thus, although both proteins localized in the cytoplasm, there were localization patterns clearly different in both CmVPS41 in some structures when expressed in *N. benthamiana* epithelial cells. From those structures, some of them (transvacuolar strands and nuclear and membrane/tonoplast spots) would be related with the susceptible variant, whereas the intravacuolar invaginations would be related with the resistant CmVPS41SC variant.

**Figure 2.**
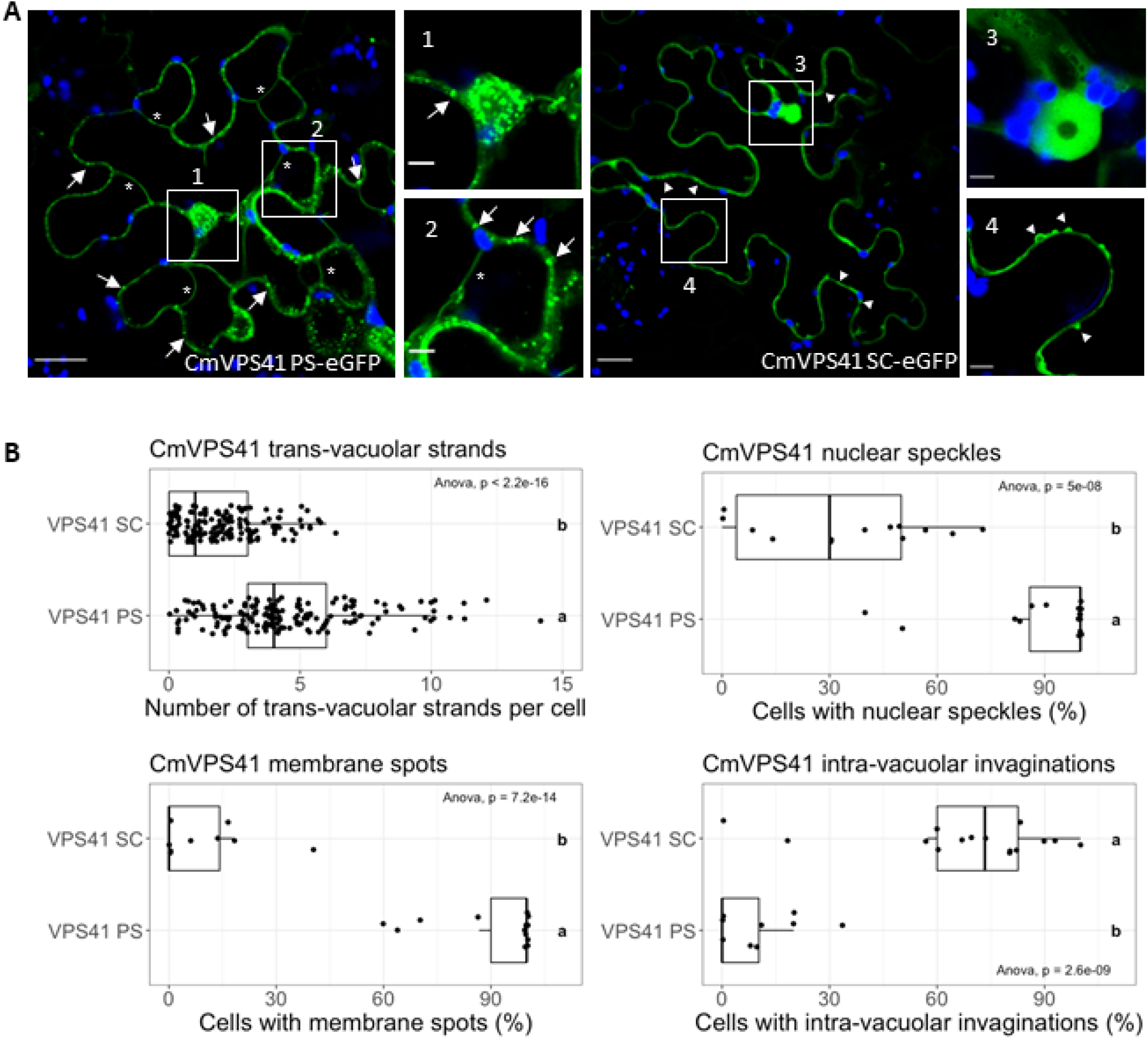
Localization patterns of CmVPS41. **A**. Localization of CmVPS41PS (CmVPS41PS-eGFP) and CmVPS41SC (CmVPS41SC-eGFP). In blue, chloroplast autofluorescence. Innage scales from whole images correspond to 20 pm and amplified images scale is 5 pm. Arrows: membrane spots. Arrowheads: intravacuolar invaginations. *: Transvacuolar strands. **B**. Boxplots of distinctive structures in CmVPS41PS and SC-expressing cells. Each boxplot was generated with R package ‘ggpubr*. Significant one-way analysis of variance (ANOVA) (p-value<0.05) between each CmVPS41 and specific structures is shown in each boxplcc Post-hoc Tukey results within treatments are indicated with letters. The same letter corresponds to nonsignificant differences between CmVPS41PS and CmVPS41SC.

**Fig 3.**
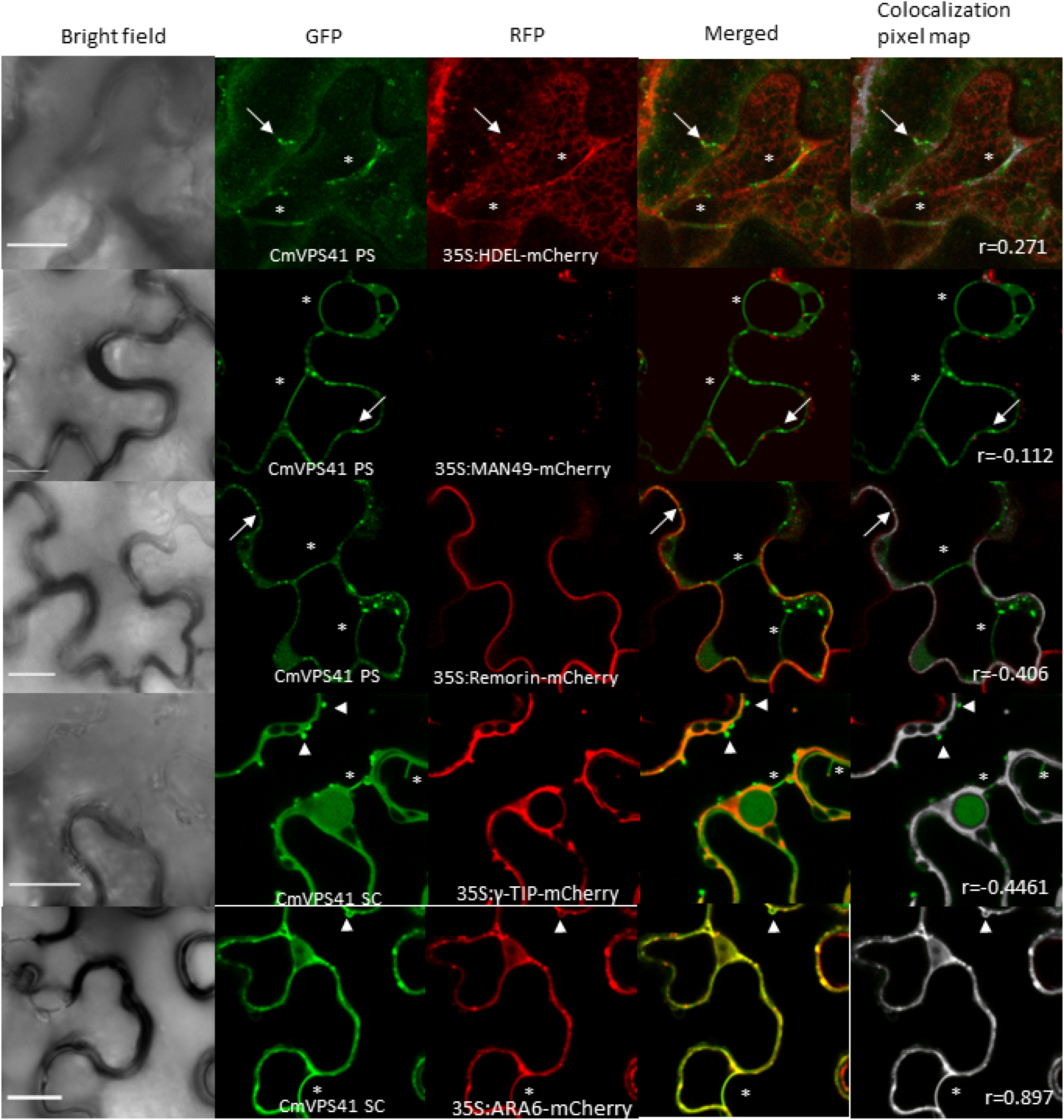
Colocalization of CmVPS41 with organelle markers. GFP channel: CmVPS41s specific structures. RFP channel: different organelle markers, endoplasmic reticulum (35S: HDEL-mCherry), Golgi apparatus (35S: MAN49-m Cherry), plasma membrane (35S: R e mo rin-m Cherry), tonoplast (35S:Y-TlP-mCherry) and late endosome {35S:ARA6-mCherry). Merged: co-localization is shown as a yellow colour. Colocalization pixel map, shown in grey, while non-colocalized pixels keep the original colour. Pearson correlation coefficient (r) of pixel matching colocalization was calculated with Fiji software analysis tool “Colocalization threshold”. Scale bars are 20 pm. Arrows indicate membrane spots. Arrowheads indicate intravacuolar invaginations. Asterisks indicate transvacuolar strands.

**Figure 4.**
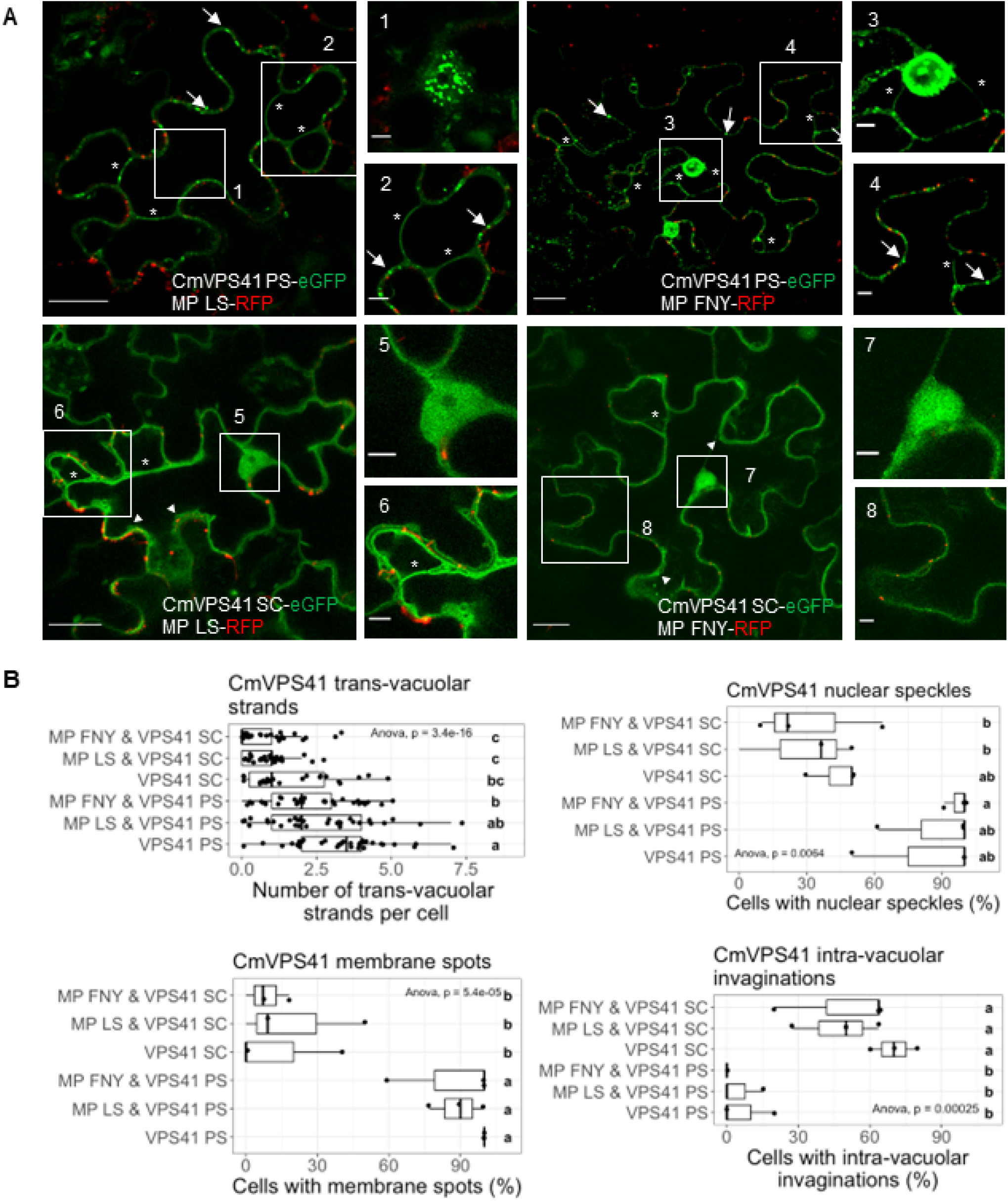
CMV MPs effect on CmVPS41-induced structures. **A**. Co-localization of CmVPS41PS-eGFP or CmVSP41SC-eGFP with CMV MP-FNY-RFP or CMV MP-LS-RFP. Numbers indicate areas amplified on the right. Inlet 1 shows the nucleus in a different Z plane than in the whole image. Scale bars are either 20 pm (whole image) or 5 pm (amplified image). Arrows indicate membrane spots. Arrowheads indicate intravacuolar invaginations. Asterisks indicate transvacuolar strands. **B**. Boxplots of CmVPS41-induced structures. Each boxplot was generated with R package ‘ggpuhr’. Significant one-way analysis of variance (ANOVA) (p-value<0.05) between each treatment and specific CmVPS41 structure is shown in each boxplot. Post-hoc Tukey results of treatments are indicated with letters. The same letter corresponds to nonsignificant differences between CmVPS41PS and CmVPS41SC(co-infiltrated or not with CMV MPs).

### The differential structures co-localize with late endosomes

VPS41 is a protein involved in the transport of cargo proteins from late Golgi to the vacuole via either vesicles or endosomes. To identify the sub-cellular nature of the distinctive structures, a set of cell markers were used. As seen in Figure 3, CmVPS41 from both variants are identified in several organelles, including endoplasmic reticulum, plasma membrane, tonoplast, or late endosomes. However, the transvacuolar strands and the spots seen in CmVPS41PS expression and the invaginations from CmVPS41SC colocalize only with late endosome (Figure 3 and Supplemental Figure S2). Controls expressing the Ara6 late endosome marker alone does not show those typical structures (Supplemental Figure S5 and S6). Thus, the differential structures shown by expression of VPS41s are derived of late endosomes, and this suggests that the expression of both variants induces a different distribution of the endosomes throughout the cytoplasm, including crossing the vacuole from side to side and invaginations towards the vacuole.

### The different distribution of late endosomes does not change with the co-expression with CMV MP

The determinant of virulence that relates to the *cmv1* gene (CmVPS41SC) is the MP (Guiu-Aragonés et al., 2015). As the MP from CMV-FNY enables the virus to overcome the resistance posed by *cmv1*, whereas the MP from CMV-LS does not, it might be possible that the localization pattern of CmVPS41SC could change in the presence of MP FNY. As shown in Figure 4, co-agroinfiltration experiments expressing both CmVPS41s and both CMV MPs showed that the localization pattern of CmVPS41PS does not change significantly in the presence of either MP, showing no significant differences in nuclear speckles, membrane spots and absence of intravacuolar invaginations. Only the presence of either MP seemed to decrease slightly the number of transvacuolar strands per cell (Figure 4B). Likewise, the localization of CmVPS41SC did not change in the presence of either MP LS or MP FNY, showing no speckles, very few transvacuolar strands and most cells had tonoplast invaginations, like the pattern of the CmVPS41SC expressed alone (Figure 4B). These experiments were repeated three times with comparable results. Therefore, the co-expression of CmVPS41SC with the MP FNY, does not induce any significant change in the localization of the CmVPS41SC that could resemble a susceptible pattern, more like that of CmVPS41PS. Additionally, Figure 4A shows that membrane spots do not co-localize with PDs, since those spots do not co-localize with MPs.

### Localization patterns of CmVPS41 from resistant and susceptible melon genotypes is different

The analysis of the CmVPS41 gene in 54 melon accessions had identified several new resistant genotypes, among them, Freeman’s Cucumber (FC) and Pat-81, (Pascual et al., 2019). CmVPS41 from SC, differ from CmVPS41PS in three amino acid positions (P262A, L348R and S620P). However, only the change L348R correlates with resistance to CMV (Giner et al., 2017; Pascual et al., 2019). Pat-81 and FC differ from PS in the same P262A, S620P and also in G85E instead of L348R (Pascual et al., 2019) (Figure 5A). On the contrary, the accession Cabo Verde is a susceptible genotype that shared P262A and S620P amino acid polymorphisms with SC, FC and Pat-81, but lacks any putative causal mutation (Figure 5A). Over-expression of CmVPS41 from both FC and Pat-81 genotypes, as well as from Cabo Verde, confirmed most of the above characteristics. CmVPS41 from Cabo Verde presented membrane spots and nuclear speckles, transvacuolar strands and very few intravacuolar invaginations, like PS does (Fig 5B, C), whereas CmVPS41 from the accessions FC and Pat-81 showed smooth nuclei and cytoplasm and very few transvacuolar strands, like CmVPS41SC. However, unlike CmVPS41SC, they show very few intravacuolar invaginations (Figure 5B, C). Thus, this suggests that the resistance to CMV would not be related to the presence of invaginations of the tonoplast. Western blot analyses showed that all proteins were expressed, although those from CV and FC in lower amount. However, independently of the amount of protein expressed, PS and CV susceptible accessions showed the same subcellular localization and the same is true for FC and Pat-81 resistant accessions (Supplemental Figure S3).

**Figure 5.**
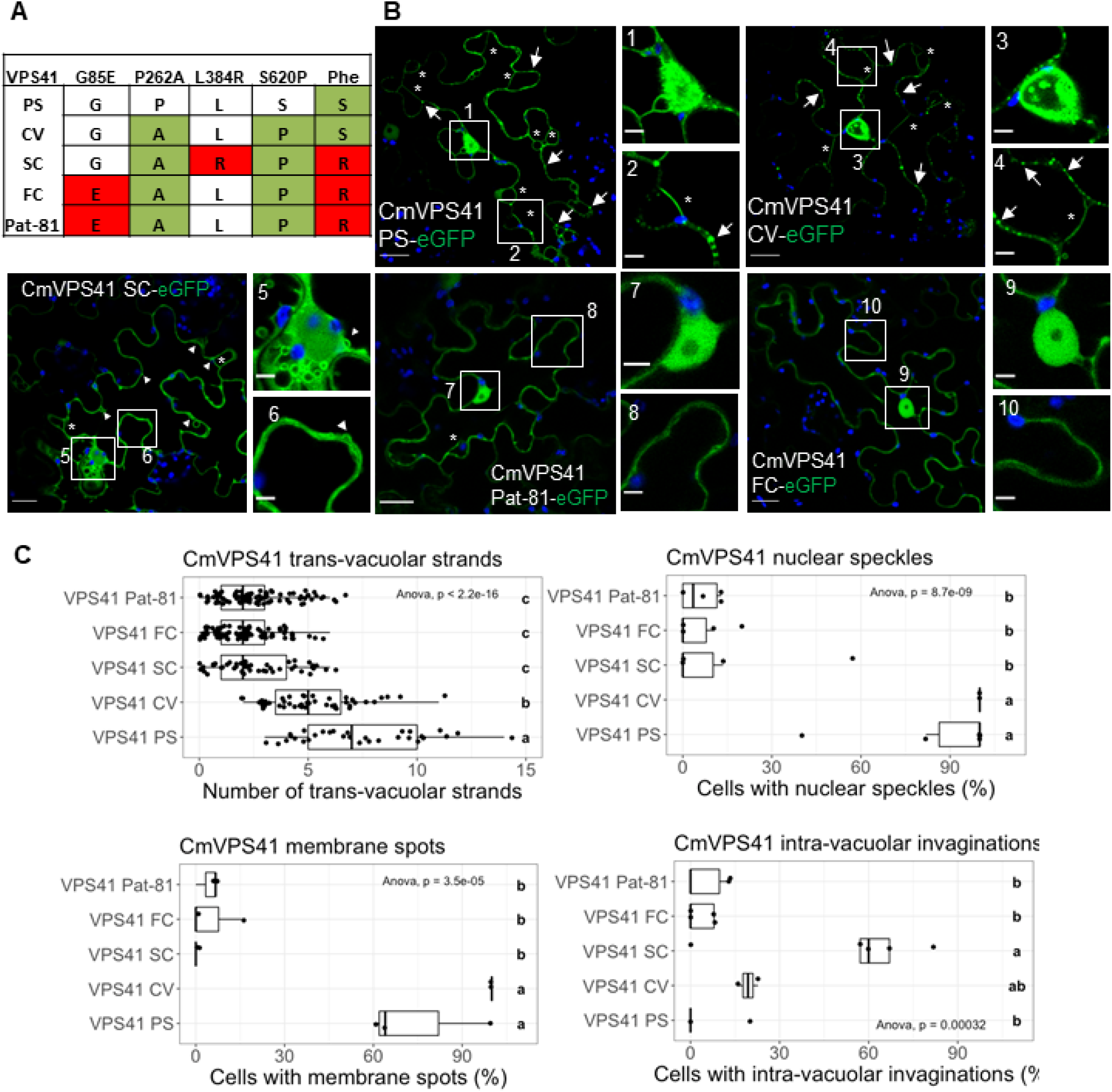
CmVPS41-induced structures in exotic melon genotypes. A. Amino acid changes of the CmVPS41 melon variante compared to CmVPS41PS. Phe: Phenotype of the corresponding variant S: Susceptible to CMV-LS. R: Resistant to CMV-LS **B**. CmVPS41 distinctive structures in the melon genotypes CV (CmVPS41CV-eGFP), P at-SI (CmVPS41Pat-81-eGFP), FC (CmVPS41FC-eGFP), PS (CmVPS41PS-eGFP) and SC (CmVPS41SC-eGFP). In blue, chloroplast autofluorescence. Scale bars are either 20 pm (whole images) or 5 pm (amplified images). Arrows indicate membrane spots. Arrowheads indicate intravacuolar invaginations. Asterisks indicate transvacuolar strands. **C**. Boxplots of CmVPS41 structures in CV, Pat-81 and FC compared to PS and SC. Each boxplot was generated with R package ‘ggpubr’. Significant one-way analysis of variance (A NO VA) (p-value<0.05) between each CmVPS41 variant and each specific structure is shown in each boxplot. Post-hoc Tukey results within CmVPS41 genotypes are indicated with letters. The same letter corresponds to nonsignificant differences between CmVPS41 genotypes.

### The causal mutations induce a decrease in the number of transvacuolar strands

To determine which of the structures were related to the causal mutations, and hence, to resistance, two new constructs were built. Both kept the sequence of CmVPS41PS, but carried only the nucleotide change either at nucleotide 254, which produced the amino acid substitution G85E, or at nucleotide 1043, which expressed a VPS41 protein carrying the L348R change (Pascual et al., 2019). After infiltration into *N. benthamiana* plants, the construct carrying L348R lost the localization pattern of CmVPS41PS and showed many intravacuolar invaginations, fewer cells with membrane spots and nuclear speckles, and fewer transvacuolar strands. This is a typical CmVPS41SC localization (Figure 6A, B). Moreover, the number of cells with intravacuolar invaginations was higher than when expressing CmVPS41SC and they appeared to be generated frequently nearby the nuclear membrane. Therefore, the same mutation responsible for the resistance correlates with changes in the distribution of the late endosome to produce numerous intravacuolar invaginations and few transvacuolar strands. The overexpression of the CmVPS41 construct carrying the G85E causal mutation showed no cells with tonoplast invaginations and very few transvacuolar strands per cell (Figure 6A, B), which confirmed the observations made with Pat-81 and FC. The only difference with those accessions was the presence of some cells with membrane spots. Western blots demonstrated that all proteins were being expressed, although the construct with the mutation L348R in lower amount (Supplemental Figure S4). However, irrespective of the amount of protein expressed, CmVPS41 L348R showed the same intracellular localization than the corresponding CmVPS41SC. Altogether, these results indicate that the lack of transvacuolar strands, rather than the presence of tonoplast invaginations, correlate with the mutations that cause resistance to CMV.

**Figure 6.**
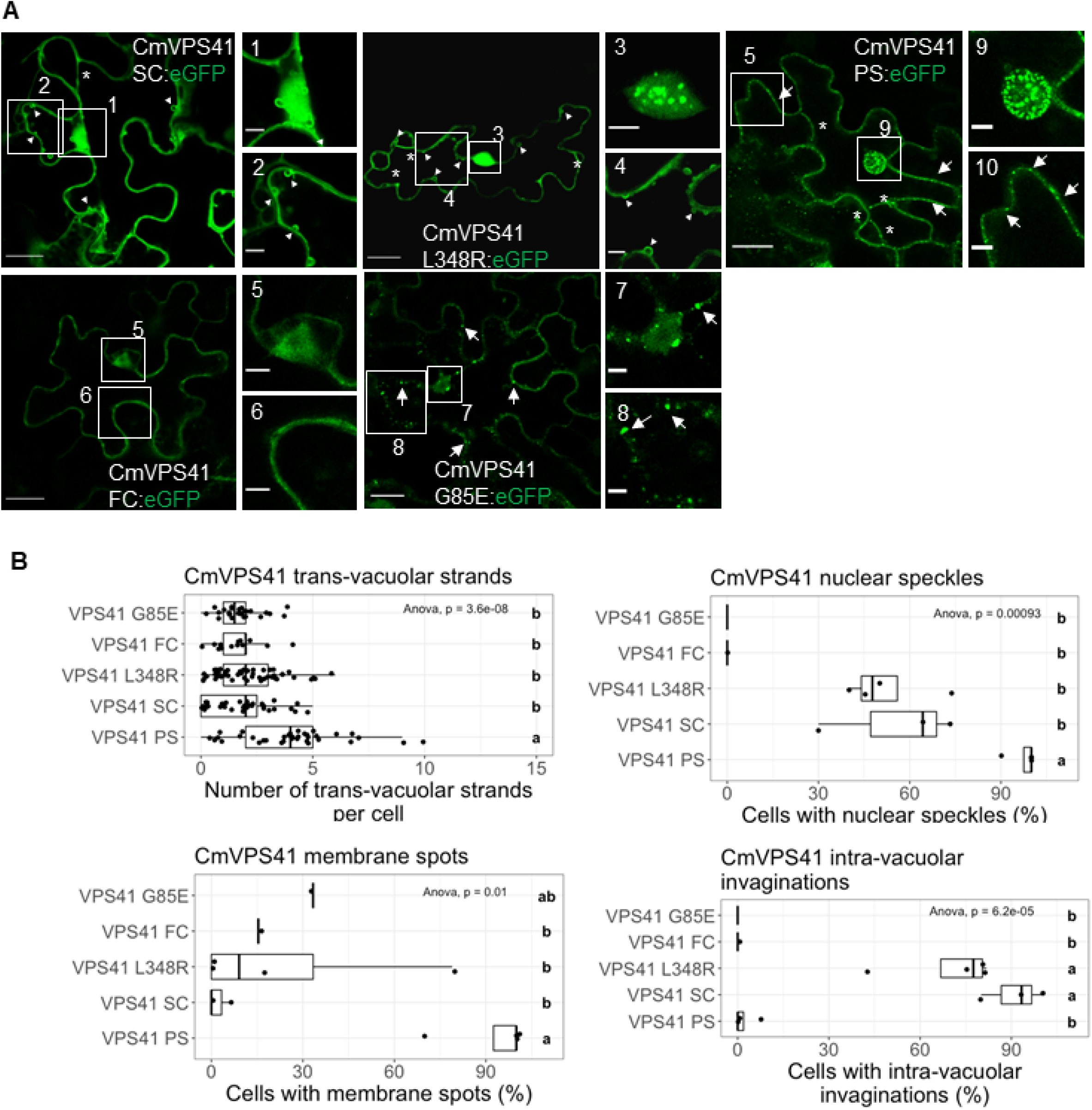
Effect of causal mutations on CmVPS41 structures. **A**. Localization of CmVPS4l carrying different causal mutations. CmVPS41 GS5E-eGFP and CmVSP41 L348R-eGFP carry the causal mutations G35E and L34BR, respectively. CmVPS41PS-eGFP_r_ CmVPS41SC-eGFP and CmVPS41FC-eGFP correspond to PS, SC and FC genotypes, respectively. Ear scales are either 20 pm (whole images) or 5 pm (amplified images). Arrows indicate membrane spots. Arrowheads indicate intravacuolar invaginations. Asterisks indicate transvacuolar strands. **B**. Boxplots of CmVPS4l structures. Each boxplot was generated with R package ‘ggpubr’. Significant one-way analysis of variance (ANQVA) (p-value<0.05) is shown in each boxplot Post-hocTukey results within treatments are indicated with letters. The same letter corresponds to no significant differences between different treatments.

### Some structures from both, resistant and susceptible CmVPS41s variants, re-localize during the viral infection

Although the MP is the virulence determinant of CMV against the gene *cmv1*, its co-expression under a strong promoter with CmVPS41 does not show an influence in the VPS41 localization pattern. However, during a real infection the MP can be expressed on due amount and time, and this could lead to a change in the CmVPS41 localization. To analyze this, *N. benthamiana* plants were inoculated either with CMV-FNY or CMV-LS and, after the onset of symptom appearance, new, symptomatic leaves were tested for virus presence by reverse transcription-PCR (Supplemental Figure 7) and then agroinfiltrated either with CmVPS41PS-GFP or CmVPS41SC-GFP. In these *N. benthamiana* infected cells, both CmVPS41PS and CmVPS41SC-infiltrated plants showed some differences in the localization pattern independently of the virus strain used. Nuclear speckles were present in many CmVPS41PS expressing cells and much fewer in CmVPS41SC-expressing cells, like in non-infected infiltrated cells shown above (Figure 7). Conversely, the number of cells with membrane spots increased, and the number of tonoplast invaginations decreased in CmVPS41SC-expressing cells with respect to non-infected plants (Figure 7A, B). Interestingly, transvacuolar strands remained very scarce for SC-expressing cells but in PS-infiltrated cells its number decreased significantly during the viral infection, suggesting that these strands could be transiently formed during a real infection and, once the infection is set, they could be no longer needed. Thus, two distinctive structures were differently present in CMV-infected cells with respect to their non-infected counterparts: the number of cells with membrane spots increased in CMV-infected CmVPS41SC-expressing cells and the number of transvacuolar strands decreased in CmVPS41PS-expressing cells.

**Figure 7.**
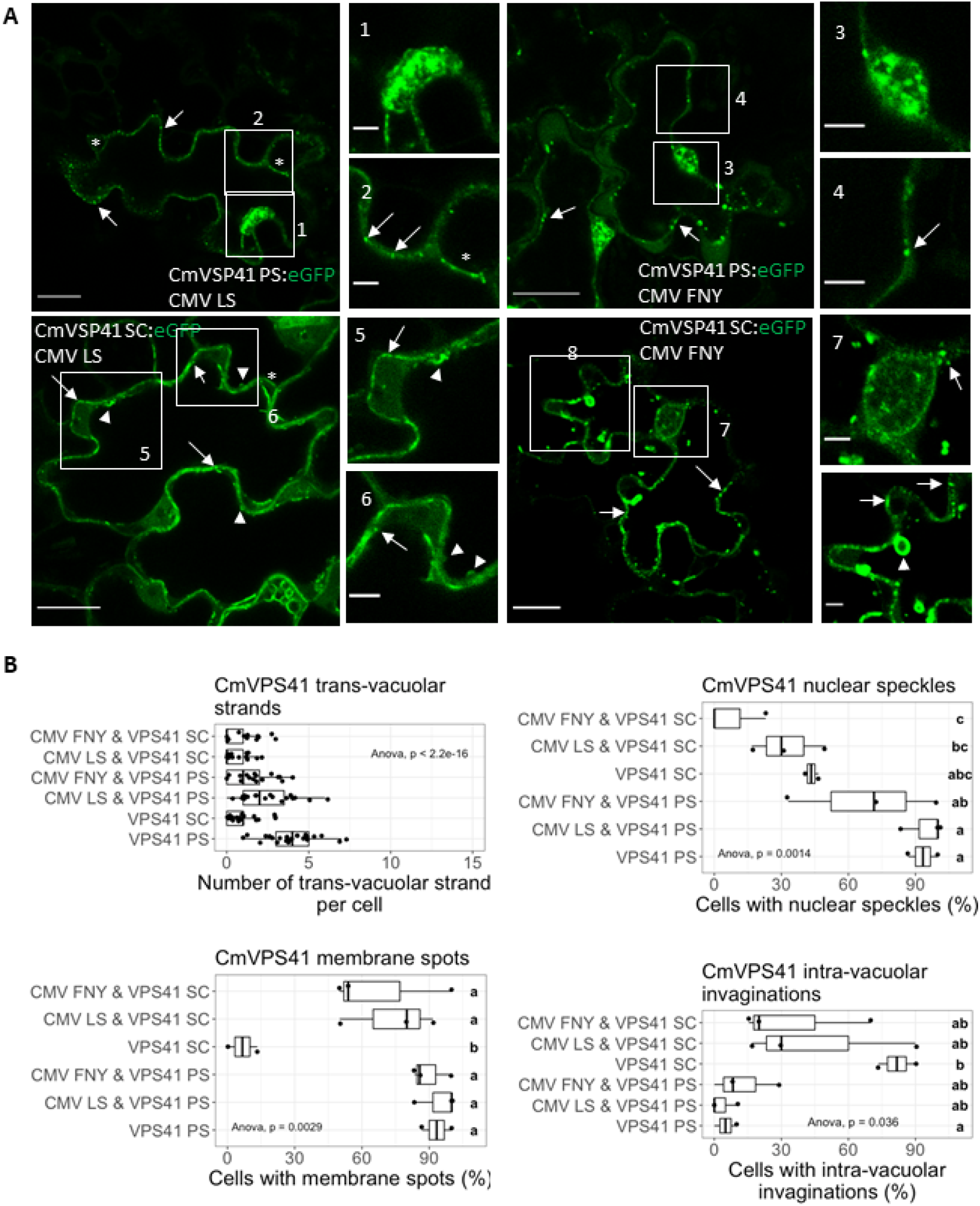
Effect of CMV infection on CmVPS41 structures. **A**. Localization of CmVPS41PS-eGFP or CmVSP41SC-eGFP in either CMV-LS or CMV-FNY infected leaves. Bar scales are either 20 pm (whole images) or 5 pm (amplified images). Arrows indicate membrane spots. Arrowheads indicate intravacuolar invaginations. Asterisks indicate transvacuolar strands. **B**. Boxplots of CmVPS41 structures. Each boxplot was generated with R package ‘ggpubr’. Significant one-way analysis of variance (ANOVA) (p-value<0.05) is shown in each boxplot Post-hoc Tukey results within treatments are indicated with letters. The same letter corresponds to non significant differences between different treatments.

## Discussion

VPS41, as a component of the HOPS complex, is a key regulator of cell trafficking. Here we have described the cellular localization of CmVPS41 both from the susceptible cultivar PS and from the resistant exotic cultivar SC, showing that they have differences, mainly in membrane and nuclear speckles, transvacuolar strands and intravacuolar invaginations. Sometimes, these structures moved during the observation. Transvacuolar strands and intra vacuolar invaginations have already been described in soybean (Nebenführ et al., 1999) and in several cell types in *A. thaliana*, where these structures are moving and changing their morphology in a manner dependent on actin microfilaments. Furthermore, the movement of the transvacuolar strands were dependent on actin microfilaments (Uemura et al., 2002). Interestingly, CMV MP has been involved in severing acting filaments to increase the size exclusion limit of plasmodesmata (Su et al., 2010). These trans vacuolar structures have been related to the distribution of different solutes, mRNAs and organelles, like Golgi vesicles in the cytoplasm up to the PDs. Moreover, the non-mobile RFP mRNA was targeted to PDs through the transvacuolar strands when co-expressed with *Tobacco mosaic virus* MP, suggesting that these strands can be used by a viral MP to direct the virus towards the PDs (Luo et al., 2018). Thus, these structures could well be used also by CMV during the infection through its MP and, together with CmVPS41, for its intracellular movement towards the PDs. The localization of CmVPS41s carrying only the causal mutations for resistance L348R (Giner et al., 2017) and G85E (Pascual et al., 2019), indicated that the resistance was mostly related to the absence of transvacuolar strands, rather than with the presence of intravacuolar invaginations, which supports the idea that those strands could be involved in CMV intracellular trafficking. A more exhaustive analysis with three-dimensional visualization of confocal stack-images shows the structure of these transvacuolar strands, appearing as ropes arranged in a fence-like manner (Figure 8). The special structures colocalize with late endosomes, which is one of the ways the CmVPS41, as part of the HOPS complex, carries cargo proteins to the vacuole. The other way, the AP-3 pathway, skips the endosome pathway and needs VPS41 to drive vesicles directly to the vacuole (Rehling et al., 1999). Thus, the late endosomal localization of the differential structures observed in our experiments suggests that CMV uses CmVPS41 in the endosomal pathway and is independent on the vesicle transport of the AP-3 pathway. Furthermore, as the co-expression with the viral MP does not have an effect in the differential structures present with CmVPS41SC, probably MP alone is not redirecting endosomal trafficking of CmVPS41 and the way in which these two proteins cooperate remains to be seen.

**Figure 8.**
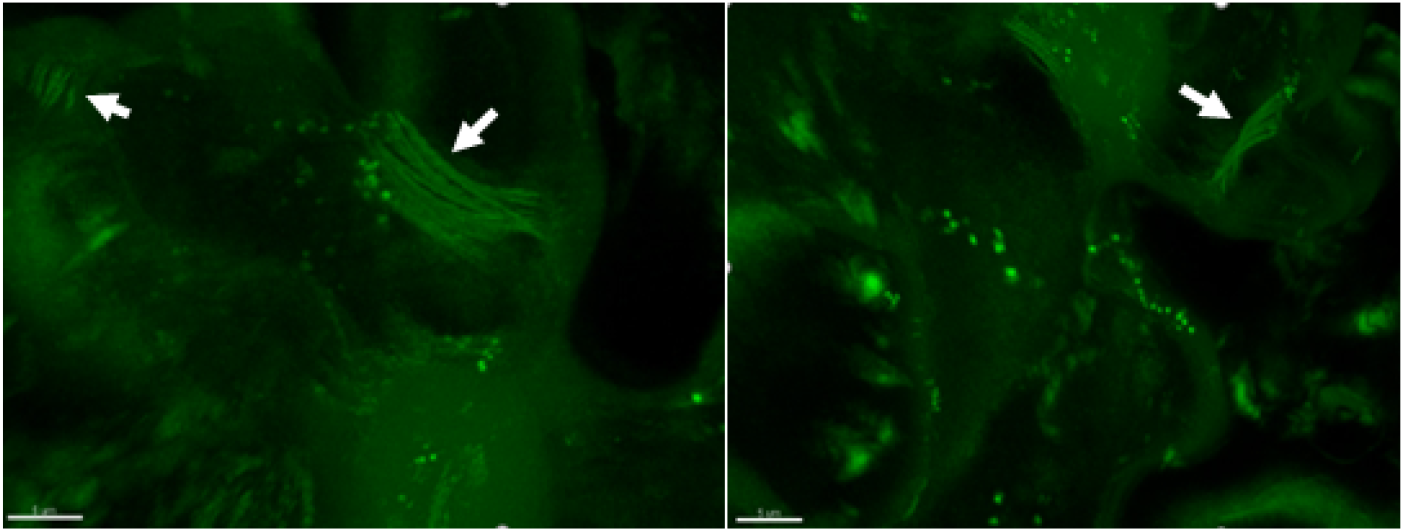
Three-dimensional reconstruction from Z-stack of CmVPS41PS-derived transvacuolar strands. Arrows indicate transvacuolar strands. N indicates nucleus. Scale bars are 5 pm.

Despite the MP being the determinant of virulence that communicates with CmVPS41 during infection, overexpression of CmVPS41s with CMV MPs does not change significantly the CmVPS41SC localization. This would suggest that either the differential structures are not really related to resistance to CMV or that the MP does not participate at this step of the infection. Alternatively, in the context of a viral infection, it may need the participation of other viral proteins to fully re-localize those differential structures. In fact, overexpression of CmVPS41s in systemically infected *N. benthamiana* leaves showed a similar localization pattern with both proteins, presenting many cells with membrane spots, very few transvacuolar strands per cell and very few cells with intravacuolar invaginations. Only nuclear speckles were behaving as in the absence of infection, abundant in PS and scarce in SC. Membrane spots do not co-localize with PDs, since they do not co-localize with MPs. Besides, membrane spots are not related to cell-to-cell viral movement, since in the resistant accessions CMV can move cell-to-cell in the inoculated leaf until reaching the bundle sheath cells (Guiu-Aragonés et al., 2016). In the context of viral infection, not only the MP is present, but all viral proteins, whose expression should be tightly regulated. For example, some MPs, accumulate transiently in the first stages of the cell infection (Maule and Palukaitis, 1991). The fact that the transvacuolar strands are dynamic (Uemura et al., 2002), suggests that, when the infection is fully established, these structures may disappear, since they may not be further needed for moving the virus towards the PDs. One possible mechanism for the dynamism of these structures during viral infection may be the timely degradation by the Ubiquitination Proteasome System (UPS). Many viral proteins are degraded *in vivo* by the UPS, among them, TMV MP as well as those MPs from *Turnip yellow mosaic virus* (TYMV), *Potato leafroll virus* (PLRV) and TGBp3 from *Potato virus X* (PVX) are known proteasome degraded viral MPs (Reichel and Beachy, 2000; Drugeon and Jupin, 2002; Vogel et al., 2007; Ju et al., 2008). Thus, UPS degradation of viral proteins, and particularly MPs, seems to be a quite extended regulation system in viral infections (for a review, see in (Alcaide-Loridan and Jupin, 2012). Thus, the interplay of all viral proteins and their timely regulation during infection could rend a CmVPS41 localization pattern quite different in different times of the infection and different than when overexpressed with MP. Further work is needed to unveil the role of CmVPS41-produced transvacuolar strands during CMV infection and the link with factors that regulate VPS41 function, such as the levels of phosphatidyl inositol or the role of Rab GTPases (Brillada et al., 2018).

## Acknowledgements

We thank Fuensanta Garcia for her technical support. This work was supported by the grants AGL2015-64625-C2-1-R and RTI2018-097665-B-C2, from the Spanish Ministry of Economy and Competitivity (cofunded by FEDER funds) and by the CERCA Programme/Generalitat de Catalunya. We acknowledge financial support from the Spanish Ministry of Science and Innovation-State Research Agency (AEI), through the “Severo Ochoa Programme for Centres of Excellence in R&D” 2016-2019 (SEV-2015-0533) and CEX2019-000902-S.

## Author contributions

N.R., I.V., I.S. and C.G-A performed research. N.R. analyzed the data and A.M.M.-H. designed research and wrote the manuscript.

## References

Alcaide-Loridan C, Jupin I (2012) Ubiquitin and plant viruses, let’s play together! Plant Physiol 160: 72–82

Amari K, Boutant E, Hofmann C, Schmitt-Keichinger C, Fernandez-Calvino L, Didier P, Lerich A, Mutterer J, Thomas CL, Heinlein M, Mély Y, Maule AJ, Ritzenthaler C (2010) A family of plasmodesmal proteins with receptor-like properties for plant viral movement proteins. PLoS Pathog 6: e1001119

Asensio CS, Sirkis DW, Maas JW, Egami K, To T-L, Brodsky FM, Shu X, Cheng Y, Edwards RH (2013) Self-assembly of VPS41 promotes sorting required for biogenesis of the regulated secretory pathway. Developmental cell 27: 425–437

Balderhaar HJk, Ungermann C (2013) CORVET and HOPS tethering complexes – coordinators of endosome and lysosome fusion. Journal of Cell Science 126: 1307–1316

Brillada C, Zheng J, Krüger F, Rovira-Diaz E, Askani JC, Schumacher K, Rojas-Pierce M (2018) Phosphoinositides control the localization of HOPS subunit VPS41, which together with VPS33 mediates vacuole fusion in plants. Proc Natl Acad Sci U S A 115: E8305–e8314

Campbell RE, Tour O, Palmer AE, Steinbach PA, Baird GS, Zacharias DA, Tsien RY (2002) A monomeric red fluorescent protein. Proc Natl Acad Sci U S A 99: 7877–7882

Carette JE, Raaben M, Wong AC, Herbert AS, Obernosterer G, Mulherkar N, Kuehne AI, Kranzusch PJ, Griffin AM, Ruthel G, Cin PD, Dye JM, Whelan SP, Chandran K, Brummelkamp TR (2011) Ebola virus entry requires the cholesterol transporter Niemann-Pick C1. Nature 477: 340–343

Cormack BP, Valdivia RH, Falkow S (1996) FACS-optimized mutants of the green fluorescent protein (GFP). Gene 173: 33–38

Drugeon G, Jupin I (2002) Stability in vitro of the 69K movement protein of Turnip yellow mosaic virus is regulated by the ubiquitin-mediated proteasome pathway. J Gen Virol 83: 3187–3197

Edwardson JR, Christie RG (1991) Cucumoviruses. In JR Edwardson, RG Christie, eds, CRC Handbook of Viruses Infecting Legumes. CRC Press, Boca Raton, FL., pp 293–319

Essafi A, Diaz-Pendon JA, Moriones E, Monforte AJ, Garcia-Mas J, Martin-Hernandez AM (2009) Dissection of the oligogenic resistance to Cucumber mosaic virus in the melon accession PI 161375. Theoretical and Applied Genetics 118: 275–284

Giner A, Pascual L, Bourgeois M, Gyetvai G, Rios P, Picó B, Troadec C, Bendahmane A, Garcia-Mas J, Martín-Hernández AM (2017) A mutation in the melon Vacuolar Protein Sorting 41 prevents systemic infection of Cucumber mosaic virus. Scientific Reports 7: 10471

Gu Y, Innes RW (2011) The KEEP ON GOING protein of Arabidopsis recruits the ENHANCED DISEASE RESISTANCE1 protein to trans-Golgi network/early endosome vesicles. Plant Physiol 155: 1827–1838

Guiu-Aragonés C, Díaz-Pendón JA, Martín-Hernández AM (2015) Four sequence positions of the movement protein of Cucumber mosaic virus determine the virulence against cmv1-mediated resistance in melon. Molecular Plant Pathology 16: 675–684

Guiu-Aragonés C, Monforte AJ, Saladié M, Corrêa RX, Garcia-Mas J, Martín-Hernández AM (2014) The complex resistance to Cucumber mosaic cucumovirus (CMV) in the melon accession PI 161375 is governed by one gene and at least two quantitative trait loci. Molecular Breeding 34: 351–362

Guiu-Aragonés C, Sánchez-Pina MA, Díaz-Pendón J, Peña EJ, Heinlein M, Martín-Hernández AM (2016) cmv1 is a gate for Cucumber mosaic virus transport from bundle sheath cells to phloem in melon. Mol. Plant Pathology 17: 973–984

Hao L, Liu J, Zhong S, Gu H, Qu LJ (2016) AtVPS41-mediated endocytic pathway is essential for pollen tube-stigma interaction in Arabidopsis. Proc Natl Acad Sci U S A 113: 6307–6312

Hashimoto M, Neriya Y, Yamaji Y, Namba S (2016) Recessive Resistance to Plant Viruses: Potential Resistance Genes Beyond Translation Initiation Factors. Frontiers in microbiology 7: 1695–1695

Hipper C, Brault V, Ziegler-Graff V, Revers F (2013) Viral and cellular factors involved in phloem transport of plant viruses. Frontiers in plant science 4: 1–18

Jiang D, He Y, Zhou X, Cao Z, Pang L, Zhong S, Jiang L, Li R (2022) Arabidopsis HOPS subunit VPS41 carries out plant-specific roles in vacuolar transport and vegetative growth. Plant Physiology 189: 1416–1434

Ju HJ, Ye CM, Verchot-Lubicz J (2008) Mutational analysis of PVX TGBp3 links subcellular accumulation and protein turnover. Virology 375: 103–117

Karchi Z, Cohen S, Govers A (1975) Inheritance of resistance to Cucumber Mosaic virus in melons. Phytopathology 65: 479–481

Karimi M, Inzé D, Depicker A (2002) GATEWAY vectors for Agrobacterium-mediated plant transformation. Trends Plant Sci 7: 193–195

Klepikova AV, Kasianov AS, Gerasimov ES, Logacheva MD, Penin AA (2016) A high resolution map of the Arabidopsis thaliana developmental transcriptome based on RNA-seq profiling. The Plant Journal 88: 1058–1070

Li B, Dong X, Li X, Chen H, Zhang H, Zheng X, Zhang Z (2018) A subunit of the HOPS endocytic tethering complex, FgVps41, is important for fungal development and plant infection in Fusarium graminearum. Environ Microbiol 20: 1436–1451

Luo KR, Huang NC, Yu TS (2018) Selective Targeting of Mobile mRNAs to Plasmodesmata for Cell-to-Cell Movement. Plant Physiol 177: 604–614

Martín-Hernández AM, Picó B (2021) Natural Resistances to Viruses in Cucurbits. Agronomy 11: 23

Maule AJ, Palukaitis P (1991) Virus movement in infected plants. Critical Reviews in Plant Sciences 9: 457–473

Nebenführ A, Gallagher LA, Dunahay TG, Frohlick JA, Mazurkiewicz AM, Meehl JB, Staehelin LA (1999) Stop-and-go movements of plant Golgi stacks are mediated by the acto-myosin system. Plant Physiol 121: 1127–1142

Niihama M, Takemoto N, Hashiguchi Y, Tasaka M, Morita MT (2009) ZIP genes encode proteininvolved inmembrane trafficking of the TGN–PVC/Vacuoles. Plant and Cell Physiology 50: 2057–2068

Pascual L, Yan J, Pujol M, Monforte AJ, Picó B, Martín-Hernández AM (2019) CmVPS41 Is a General Gatekeeper for Resistance to Cucumber Mosaic Virus Phloem Entry in Melon. Frontiers in Plant Science 10

Price A, Seals D, Wickner W, Ungermann C (2000) The Docking Stage of Yeast Vacuole Fusion Requires the Transfer of Proteins from a Cis-Snare Complex to a Rab/Ypt Protein. Journal of Cell Biology 148: 1231–1238

Rehling P, Darsow T, Katzmann DJ, Emr SD (1999) Formation of AP-3 transport intermediates requires Vps41 function. Nat Cell Biol 1: 346–353

Reichel C, Beachy RN (2000) Degradation of tobacco mosaic virus movement protein by the 26S proteasome. J Virol 74: 3330–3337

Roossinck MJ (2001) Cucumber mosaic virus, a model for RNA virus evolution. Molecular Plant Pathology 2: 59–63

Sanderson LE, Lanko K, Alsagob M, Almass R, Al-Ahmadi N, Najafi M, Al-Muhaizea MA, Alzaidan H, AlDhalaan H, Perenthaler E, van der Linde HC, Nikoncuk A, Kühn NA, Antony D, Owaidah TM, Raskin S, Vieira L, Mombach R, Ahangari N, Silveira TRD, Ameziane N, Rolfs A, Alharbi A, Sabbagh RM, AlAhmadi K, Alawam B, Ghebeh H, AlHargan A, Albader AA, Binhumaid FS, Goljan E, Monies D, Mustafa OM, Aldosary M, AlBakheet A, Alyounes B, Almutairi F, Al-Odaib A, Aksoy DB, Basak AN, Palvadeau R, Trabzuni D, Rosenfeld JA, Karimiani EG, Meyer BF, Karakas B, Al-Mohanna F, Arold ST, Colak D, Maroofian R, Houlden H, Bertoli-Avella AM, Schmidts M, Barakat TS, van Ham TJ, Kaya N (2021) Bi-allelic variants in HOPS complex subunit VPS41 cause cerebellar ataxia and abnormal membrane trafficking. Brain 144: 769–780

Schoppe J, Mari M, Yavavli E, Auffarth K, Cabrera M, Walter S, Fröhlich F, Ungermann C (2020) AP-3 vesicle uncoating occurs after HOPS-dependent vacuole tethering. Embo j 39: e105117

Serrano I, Buscaill P, Audran C, Pouzet C, Jauneau A, Rivas S (2016) A non canonical subtilase attenuates the transcriptional activation of defence responses in Arabidopsis thaliana. Elife 5

Su S, Liu Z, Chen C, Zhang Y, Wang X, Zhu L, Miao L, Wang XC, Yuan M (2010) Cucumber mosaic virus movement protein severs actin filaments to increase the plasmodesmal size exclusion limit in tobacco. Plant Cell 22: 1373–1387

Uemura T, Yoshimura SH, Takeyasu K, Sato MH (2002) Vacuolar membrane dynamics revealed by GFP-AtVam3 fusion protein. Genes Cells 7: 743–753

Vinatzer BA, Teitzel GM, Lee MW, Jelenska J, Hotton S, Fairfax K, Jenrette J, Greenberg JT (2006) The type III effector repertoire of Pseudomonas syringae pv. syringae B728a and its role in survival and disease on host and non-host plants. Mol Microbiol 62: 26–44

Vogel F, Hofius D, Sonnewald U (2007) Intracellular trafficking of Potato leafroll virus movement protein in transgenic Arabidopsis. Traffic 8: 1205–1214

